# An open source platform for analyzing and sharing worm behavior data

**DOI:** 10.1101/377960

**Authors:** Avelino Javer, Michael Currie, Chee Wai Lee, Jim Hokanson, Kezhi Li, Céline N Martineau, Eviatar Yemini, Laura J Grundy, Chris Li, QueeLim Ch’ng, William R Schafer, Ellen AA Nollen, Rex Kerr, André EX Brown

**Affiliations:** MRC London Institute of Medical Sciences, London, UK; institute of Clinical Sciences, Imperial College London, London, UK; OpenWorm Foundation, San Diego, USA; Department of Biomedical Engineering, Duke University, Durham, USA; European Research Institute for the Biology of Ageing, University of Groningen, Groningen, NL; Department of Biological Sciences, Columbia University, New York, USA; MRC Laboratory of Molecular Biology, Cambridge, UK; Department of Biology, City College of the City University of New York, New York, USA; Centre for Developmental Neurobiology, King’s College London, London, UK; Calico Life Sciences LLC, South San Francisco, USA

## Abstract

Animal behavior is increasingly being recorded in systematic imaging studies that generate large data sets. To maximize the usefulness of these data there is a need for improved resources for analyzing and sharing behavior data that will encourage re-analysis and method development by computational scientists^1^. However, unlike genomic or protein structural data, there are no widely used standards for behavior data. It is therefore desirable to make the data available in a relatively raw form so that different investigators can use their own representations and derive their own features. For computational ethology to approach the level of maturity of other areas of bioinformatics, we need to address at least three challenges: storing and accessing video files, defining flexible data formats to facilitate data sharing, and making software to read, write, browse, and analyze the data. We have developed an open resource to begin addressing these challenges using worm tracking as a model.

---

To store video files and the associated feature and metadata, we use a Zenodo.org community (an open-access repository for data) that provides durable storage, citability, and supports contributions from other groups. We have also developed a web interface that enables filtering based on feature histograms that can return, for example, fast and curved worms in addition to more standard searches for particular strains or genotypes (Fig. 1 and http://movement.openworm.org/). The database consists of 14,874 single-worm tracking experiments representing 386 genotypes (building on 9,203 experiments and 305 genotypes in a previous publication^2^) and includes data from several larval stages as well as ageing data consisting of over 2,700 videos of animals tracked daily from the L4 stage to death. Full resolution videos are available in HDF5 containers that include gzip-compressed video frames, timestamps, worm outline and midline, feature data, and experiment metadata. HDF5 files are compatible with multiple languages including MATLAB, R, Python, and C. We have also developed an HDF5 video reader that allows video playback with adjustable speed and zoom (important when reviewing high-resolution, multi-worm tracking data), as well as toggling of worm segmentation over the original video to verify segmentation accuracy during playback.

**Fig 1:**
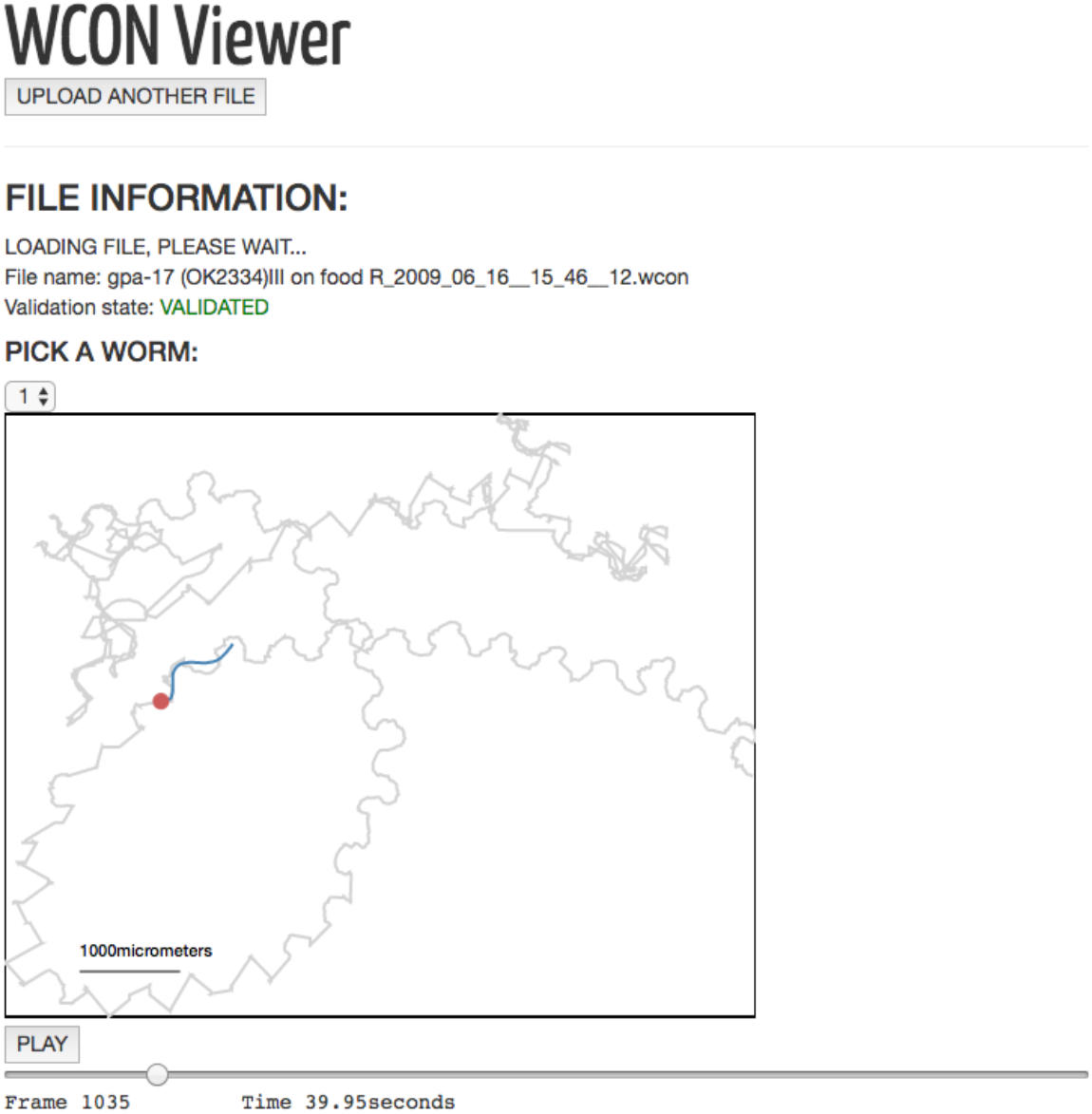
(**a**) To facilitate the sharing of tracking data, the OpenWorm Movement Database provides a web interface to search a database of worm videos by genotype, strain, and/or other discrete values. The interface includes interactive histograms, with sliders permitting users to filter results based on feature values, making it possible to find files with worms that are, for example, both fast and highly curved. The interface then connects to the video and feature data stored on Zenodo. Once downloaded, the video and feature data can be further analyzed or combined with data collected using other worm trackers through the Worm tracker Commons Object Notation (WCON), a human and machine readable JSON format for sharing tracking data. (**b**) Open-source repositories for tracking and analysis. Tierpsy Tracker segments and tracks worms, extracting the outline and skeleton of each animal then determining the head-tail orientation (Tierpsy is short for Tierpsychology, the German word for ethology). These data are saved in WCON. The Open Worm Analysis Toolbox is then used to extract the large set of behavioral features defined in Yemini *et al*.^2^. WCON includes a flexible notation for adding custom features to the WCON file if desired.

Secondly, we have defined an interchange format named Worm tracker Commons Object Notation (WCON), to facilitate data sharing and software reuse among groups working on worm behavior. WCON uses the widely supported JSON format to store tracking data as text that is both human and machine readable. It is compatible with single and multi-worm^3^ data, at any resolution: from a single point representing worm position over time^4^, to many points representing the high-resolution skeleton of a moving worm^2^. Importantly, it also supports custom feature additions so that individual labs can store their own specific data sets alongside the universal set of basic worm data. WCON readers are available for Python, MATLAB, Scala, and C. Detailed documentation for the file formats and software is available on the project page (https://github.com/openworm/tracker-commons).

Finally, we have complemented the database and file formats with open-source software written in Python for single and multi-worm tracking, feature extraction, review, and analysis (Supplementary Discussion).

The suite of tools we have reported makes quantitative behavior (re-)analysis more accessible for both experimentalists and computational scientists. It may also serve as a template for similar efforts in other model organism communities.

## Acknowledgements

This work was supported by the MRC through grant MC-A658-5TY30 to AEXB. QC is supported by an ERC Starting Grant (NeuroAge 242666), Research Councils UK Fellowship, and the University of London Central Research Fund. Some strains were provided by the CGC, which is funded by the NIH Office of Research Infrastructure Programs (P40 OD010440).

## Supplementary Discussion

### Submitting Data to movement.openworm.org

The OpenWorm movement database is intended to be a growing resource to compile and compare behavior data contributed by the community. Despite the variety of behavioral experiments, the WCON format enables multiple labs to contribute data in a form that is easily validated and from which standard behavioral features can be derived. To make WCON easier to work with we have made a web browser-based viewer that checks that a WCON file has the correct format, renders the data as a video, and displays the metadata and units in a table (Fig. 1). Once data have been validated using either the viewer or one of the WCON readers on the Worm tracker Commons project page, they can be submitted through a web form on the movement database site. Submitted data will be reviewed manually to ensure they contain worm behavior data and analyzed to extract the same features used to search the movement database. Finally, the WCON and feature data will be uploaded to the OpenWorm Movement Database community on Zenodo.org for long term storage and citability.

**Fig 1:**
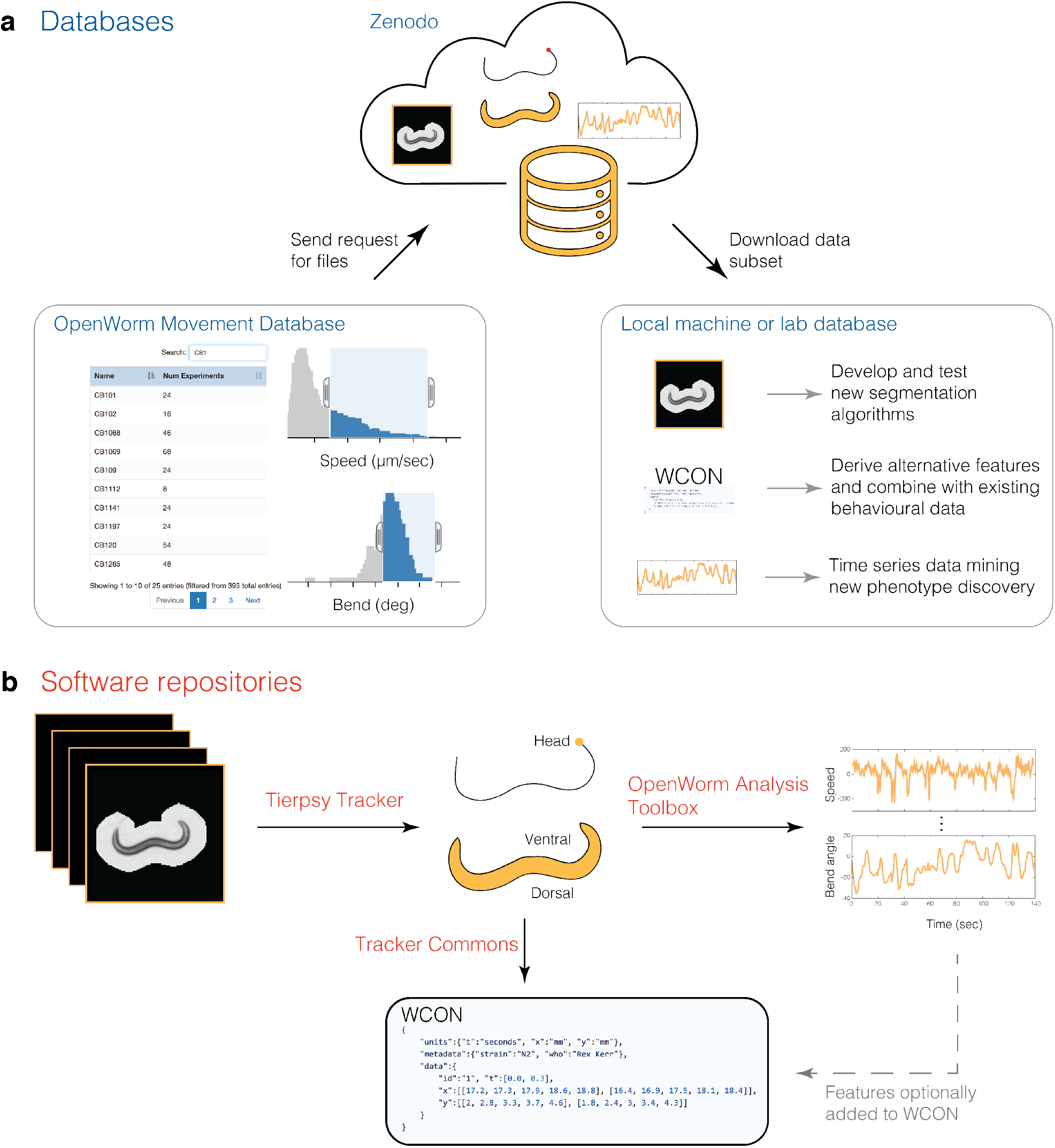
WCON viewer. Screenshot showing a validated WCON file being viewed in a web browser. The video can be played and paused, scrolled through using the slider, and zoomed. Below the video, the browser displays the metadata and units in a table.

To maximize comparability between submitted data and the existing data on http://movement.openworm.org/ we recommend the use of the same protocol that was used to collect the original data^1^. However, we recognize that experiments with different goals may require different protocols and emphasize that we accept data collected using other protocols.

### Tierpsy Tracker Description

Tierpsy Tracker is a multi-worm tracker written primarily in Python that is capable of extracting postural information. Tierpsy Tracker was designed to occupy a niche that was not filled by existing worm trackers (see Comparison with other trackers below for details). It extracts the same high-dimensional feature set as WormTracker 2.0 so that its output can be directly compared to the growing worm behavior database described here. It is fully open source, including all dependencies, so no commercial software (such as MATLAB or Labview) is required to inspect it, run it, or modify it, while executable versions are provided for both Windows and Mac OSX for those who want to use the software without dealing with the source code. It has the following main features:

- Following segmentation, it saves output videos in a compressed HDF5 format that preserves full pixel information around worms losslessly while zeroing background pixels (see Figure 1 in main text). To monitor slowly changing background features such as food depletion, full resolution images can be saved with adjustable frequency. The HDF5 files can be read using many languages (including MATLAB and Python) and support precise frame indexing. They also include experiment and analysis metadata.
- It supports a variety of video formats and experimental setups (see Sample Analysis below).
- It provides graphical user interfaces to calibrate the parameters for a new experimental setup, review the segmentation results, and manually join trajectory fragments if desired.
- It can analyze large data sets from screening projects consisting of thousands of videos by taking advantage of multicore processors to implement ‘embarrassingly parallel’ processing of multiple videos. This can be done using a simple batch processing function in the graphical user interface or from the command line.
- Its processing pipeline is modular so that analysis steps can be skipped or added depending on the analysis type and it records the provenance of output files (including software version and analysis parameters) to improve reproducibility.
- It is able to copy files from and to temporary directories prior to and after the analysis in order to deal with unstable remote connections.

### Comparison with Worm Tracker 2.0

Tierpsy Tracker is at its core a multi-worm generalization of WormTracker 2.0^1^, a single-worm tracker, although it should be emphasized that the underlying software has been ported to Python from MATLAB and completely re-designed. Tierpsy Tracker is fully compatible with videos produced by the WormTracker 2.0 hardware. The skeletonization, stage alignment, and feature calculation algorithms are Python ports of the original MATLAB algorithms. When WormTracker 2.0 files are analyzed in Tierpsy Tracker, the HDF5 output files include stage movement and experiment information along with the segmented video, reducing the risk of misplacing or losing this essential metadata.

Tierpsy Tracker uses a locally calculated threshold as opposed to the global threshold used in WormTracker 2.0, which makes it more robust to non-uniform lighting. However, on the high-contrast videos produced typically produced by WormTracker 2.0, the results are similar (see Fig. 2A). In cases where there are substantial differences between the two trackers, these are most-often caused by head-tail identification errors that are corrected by Tierpsy Tracker’s more accurate head tail detection algorithm (see “Head/tail identification”). We measured the head/tail orientation accuracy by randomly selecting 100 videos from the database and manually assessing if the orientation was correct in each frame that was successfully skeletonized. In a random subset of 100 videos, containing 1.9 x 10^6^ frames that were successfully skeletonized in both Tierpsy and WT2.0, we confirmed by manual inspection that Tierpsy made a mistake in only 216 of the frames (0.01%) while WormTracker2.0 made a mistake in 87,751 of the frames (4.49%). To further test the accuracy, we measure the RMSE between skeletons produced by WormT racker 2.0 and Tierpsy T racker in 1.8 x 10^8^ frames across 9343 videos. We consider that two skeletons are in agreement if the RMSE between them is less than 1/48 the length of the Tierpsy Tracker skeleton which is equivalent to a single segment in the skeleton. We found that 96.19% of the skeletons are in agreement, and that 8410/9343 of the videos have 90% or more of their skeletons in agreement (Fig. 2). Head-tail errors will result in a large RMSE for otherwise well-skeletonized worms so we switched the orientation of the WormTracker 2.0 skeletons and re-calculated the RMSE. If we use the lower of the two RMSE values between switched or not switched, we find that 99.20% of the skeletons are in agreement, and that 9208/9343 of the videos have 90% or more of their skeletons in agreement.

**Fig 2:**
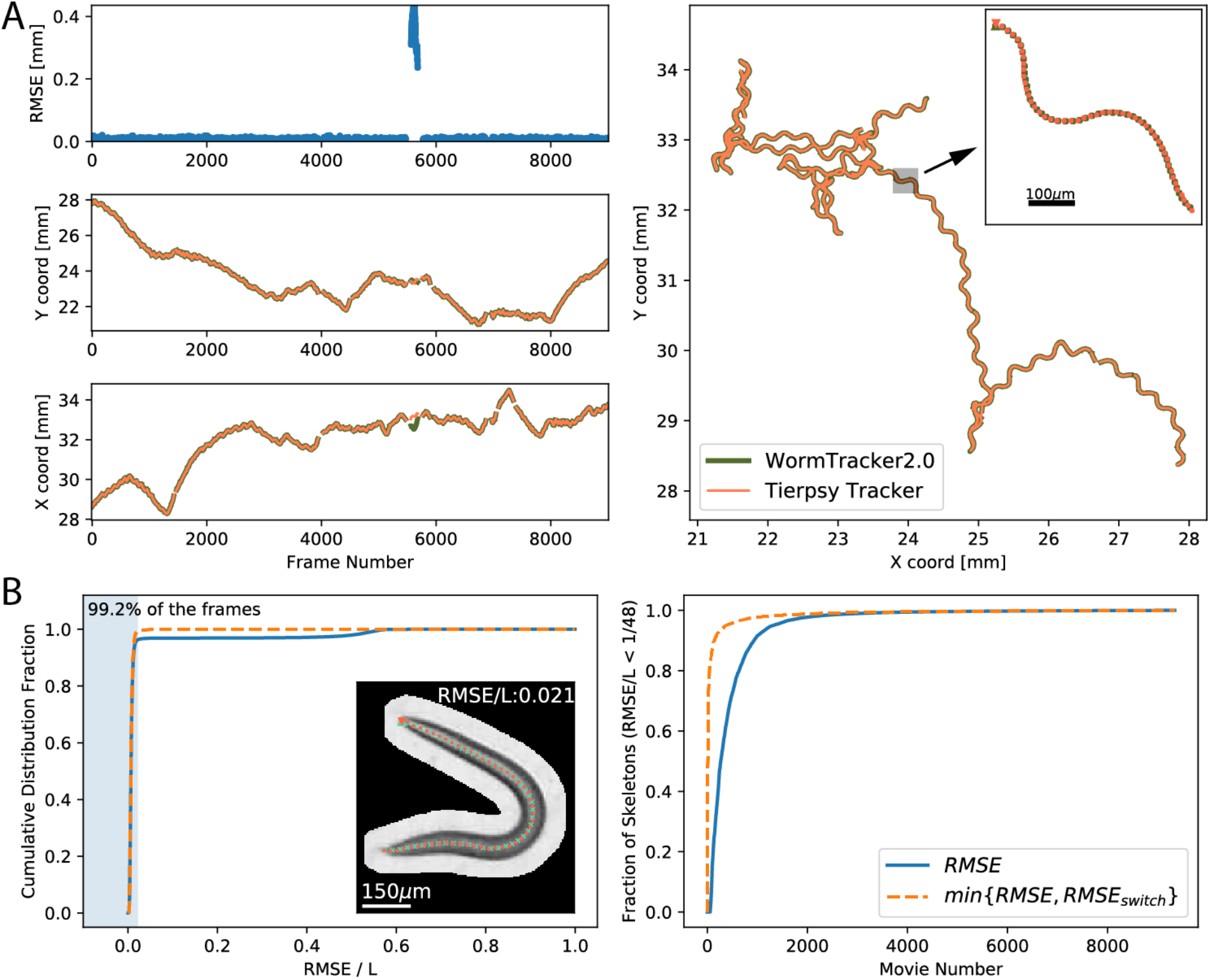
Benchmark between Tierpsy Tracker and Worm Tracker 2.0. A. Tracking results from video of a single L4 N2 analyzed with both Tierpsy Tracker (orange) and Worm Tracker 2.0 (green). Both trackers produce almost identical results except in a small segment where there was a head-tail identification error by Worm Tracker 2.0 (frames 5564 to 5678). The Tierpsy Tracker skeleton coordinates were rotated to align them with the Worm Tracker results prior plotting. This is because the coordinate system in Tierpsy matches the camera orientation, while in WT2.0, it matches the stage position. *Left top*, Root Square Mean Error (RMSE) between the skeletons output by each tracker. *Left centre and bottom*, y and x head coordinates over time. *Right*, midbody trajectory in space. *Inset*, example of the skeletonization results at frame 2500. B. *Left*, Cumulative distribution of the total fraction of skeletons with a given RMSE between the results of Worm Tracker 2.0 and Tierpsy Tracker. RMSE values are normalized by the worm length (L). The shaded blue region marks the threshold of 1/48 ≈ 0.021 and contains 99.2% of all the frames in RMSE_switch_. The inset shows an example of WT2.0 (red) and Tierpsy (cyan) with a RMSE of 0.02, which is at the border of the shaded region. 99.2% of skeleton agreements are as good or better than this. *Right*, videos sorted by the fraction of RMSE/L less than 1/48. See main text for more information.

### Comparison with other trackers

Several worm trackers have been previously described^1–19^. Most of the trackers either extract detailed postural information of one worm at a time or track multiple worms but extract only coarser-grained features such as centroid speed or orientation. Collecting postural information in addition to overall motion can be useful for phenotyping^1,13,20–22^, while tracking multiple worms can increase throughput and make it possible to study worm-worm interactions. Tierpsy Tracker is one of a relatively smaller number of trackers that extracts postural information from frames containing multiple worms and is the only tracker that compresses videos by background subtraction while maintaining uncompressed pixel information around tracked objects. This is an important feature because it makes Tierpsy Tracker compatible with large scale screens while maintaining enough information to re-analyze datasets using algorithms that depend on pixel information (e.g. egg laying detection, texture-based analyses, idTracker^23^, etc.).

There are two other published open source trackers that track multiple worms and extract detailed postural information: the Multi-Worm Tracker^8^ and CeLeST^6^. The Multi-Worm Tracker can be used for high-throughput screens but compresses data by saving worm positions and outlines while discarding all pixel information. CeLeST extracts features from videos but does not compress videos. Furthermore, Tierpsy Tracker is fully open source, including its dependencies whereas the Multi-Worm Tracker and CeLEST use commercial software for at least part of their pipelines (the Multi-Worm Tracker requires a Labview runtime license for its user interface and CeLEST is written in MATLAB). Tierpsy Tracker occupies an intermediate niche between these two other trackers in terms of processing speed and ease of use. It is slower than the Multi-Worm Tracker but includes graphical user interfaces for parameter tuning and batch processing making it more accessible for users unfamiliar with running software from the command line (although Tierpsy Tracker can also be run from the command line). Compared to CeLeST, Tierpsy tracker extracts a wider range of features (provided sufficient video resolution, Fig. 3) and includes simple parallelization without user intervention, which is convenient for running on larger datasets. Tierpsy tracker is also faster on a per-file basis: on a MacBook Pro (2.7GHz Core Quad, 16GB) the Dataset S1 in Restif *et al*.^6^ was analyzed in 3 minutes using CeLEST (MATLAB 2014b) and 1.5 minutes using Tierpsy.

**Fig. 3:**
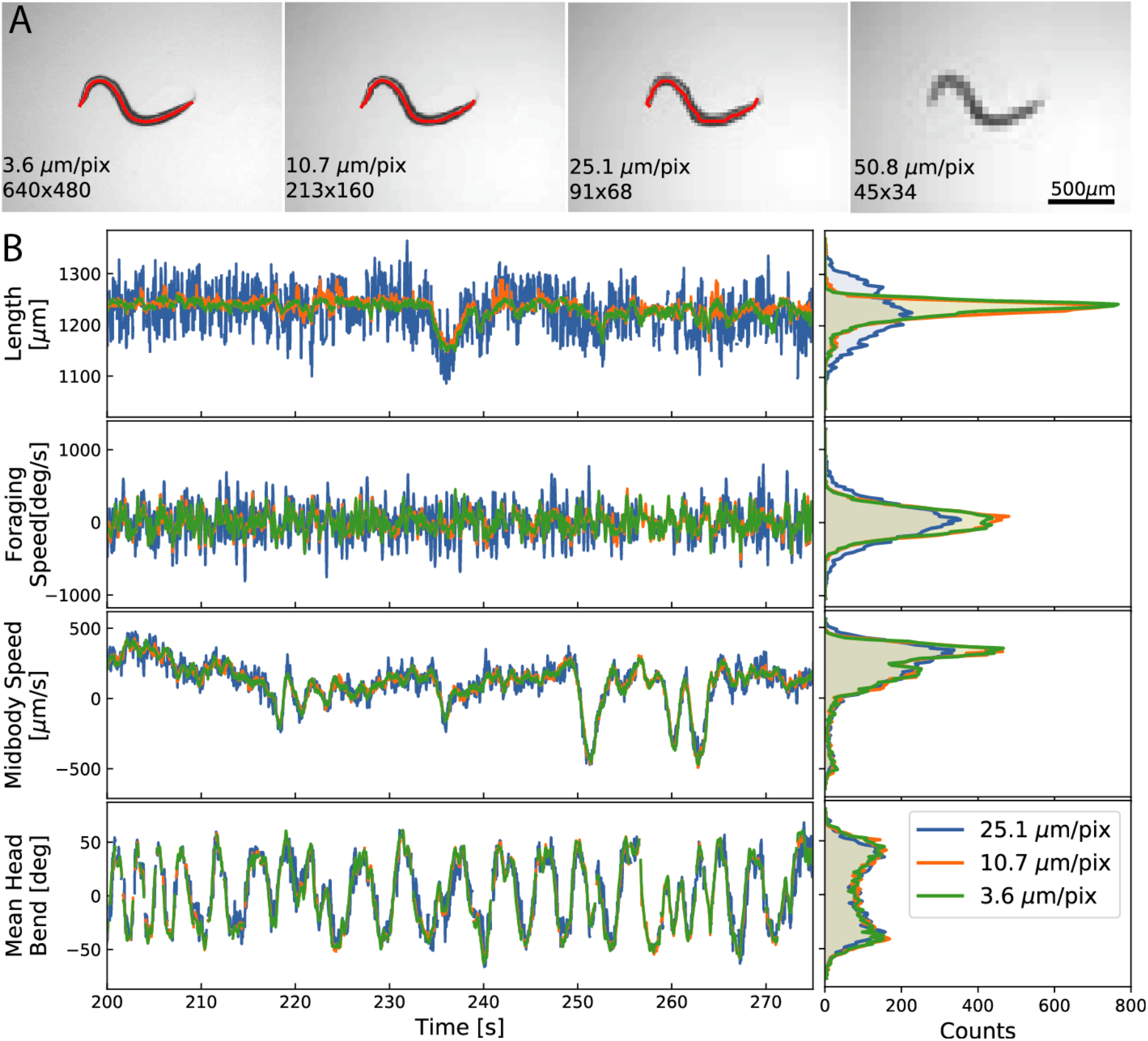
Effect of resolution on tracking. To simulate the effect of pixel size in the tracking analysis we reduced the size of a video of an N2 adult recorded using Worm Tracker 2.0. A. Example of a frame at the different image sizes tested, the skeletonization results are overlaid (red). We test image sizes of 680×480, 213×160, 91×68 and 45×34, this is the equivalent of 3.57, 10.73, 25.12 and 50.80 μm/pixel respectively. The skeletonization algorithm fails at 50.8 μm/pixel. B. Time series plots (*left*) and histograms (*right*) of a few selected features. Most of the features calculated produce similar results at different resolutions, except for features related with morphology (length, width, area) or to the head and the tail (foraging, angular speed) which show an increase in noise with reducing resolution.

### Video compression in large fields of view

An uncompressed video stream from a four-megapixel camera recording at 25fps will fill a 2TB disk in less than 3hrs of recording. Some form of compression is therefore an important component of a high-resolution tracker intended for even moderate throughput experiments.

The most common approach is to use a video format that uses lossy compression to manage file sizes while maintaining visual features tuned to human vision (although not necessarily optimal for computer vision^24^) but this introduces compression artefacts.

A more extreme approach is adopted by the Multi-Worm tracker^8^, which saves a compact set of features including worm contour and position. This greatly reduces file size, but it comes at the expense of losing all the video textural information, precluding re-analysis with improved computer vision methods. Tierpsy Tracker uses an intermediate approach by only keeping the pixels around candidate worm regions and setting the rest of the image to zero (see Compression/object detection). These segmented images are highly compressible using standard lossless compression algorithms thus reducing file sizes while maintaining full resolution information around each worm so that tracking and analysis can be repeated as improved algorithms are developed. Tierpsy Tracker also uses pixel intensities in its improved head-tail detection algorithm (see “Head/tail identification” and Fig. 4).

**Fig 4:**
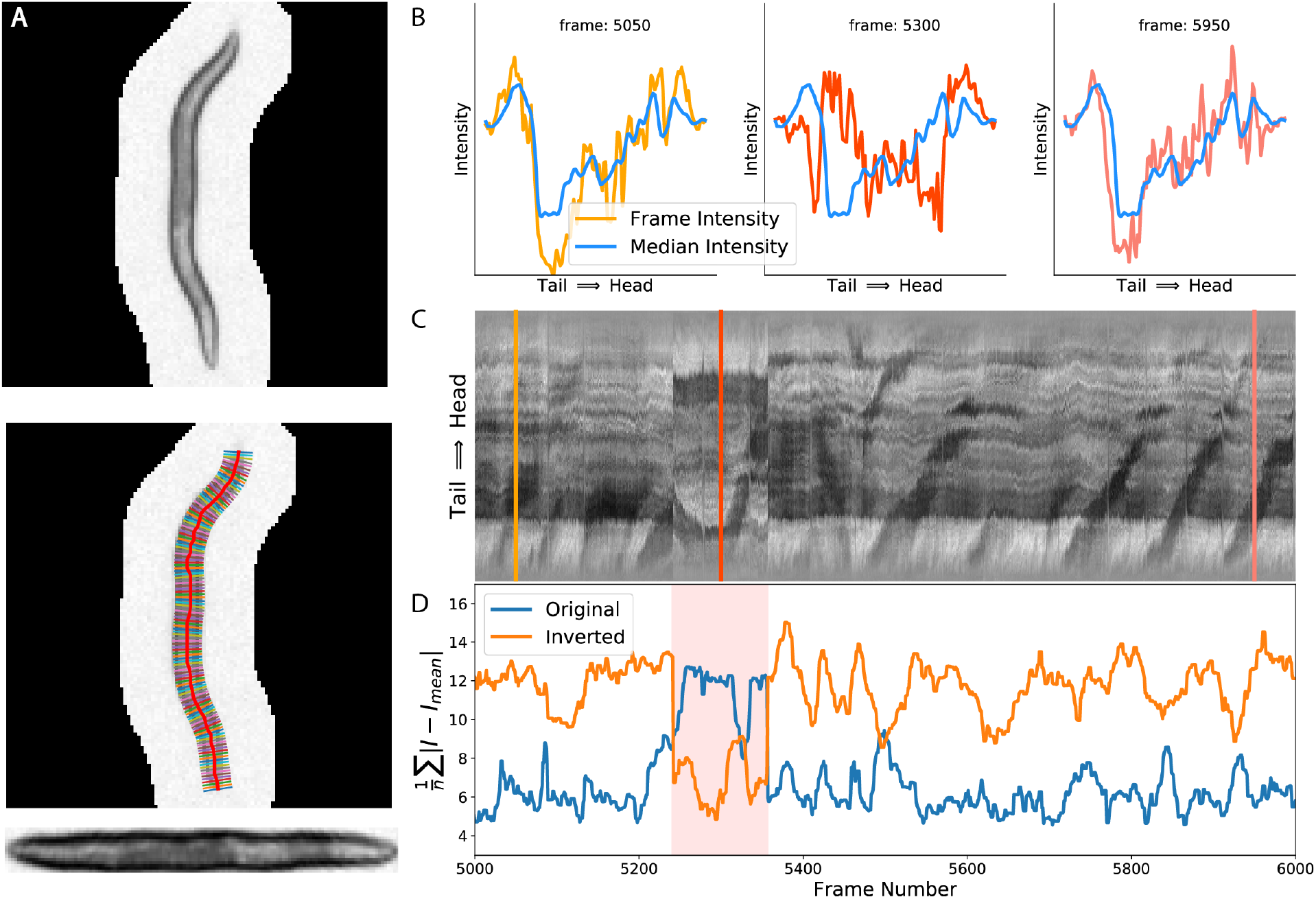
Head tail correction using intensity. A. Process of interpolation along the segments perpendicular to the worm skeletons to obtain the intensity along a straightened worm^25^. B. Example of different intensity profiles along the worm skeletons for different frames. The blue line corresponds to the median values among all the intensity profiles (global intensity profile). C. Kymograph of the intensity profile along the skeleton. The position of each the intensity profile shown in panel B is marked with the corresponding colour line. D. L1-norm between the difference of each frame intensity profile and the global intensity profile (original, blue or inverted, orange). The region corresponding to a wrongly oriented skeleton block (pink shade) has a smaller value in the inverted than in the original L1-norms.

**Fig. 5:**
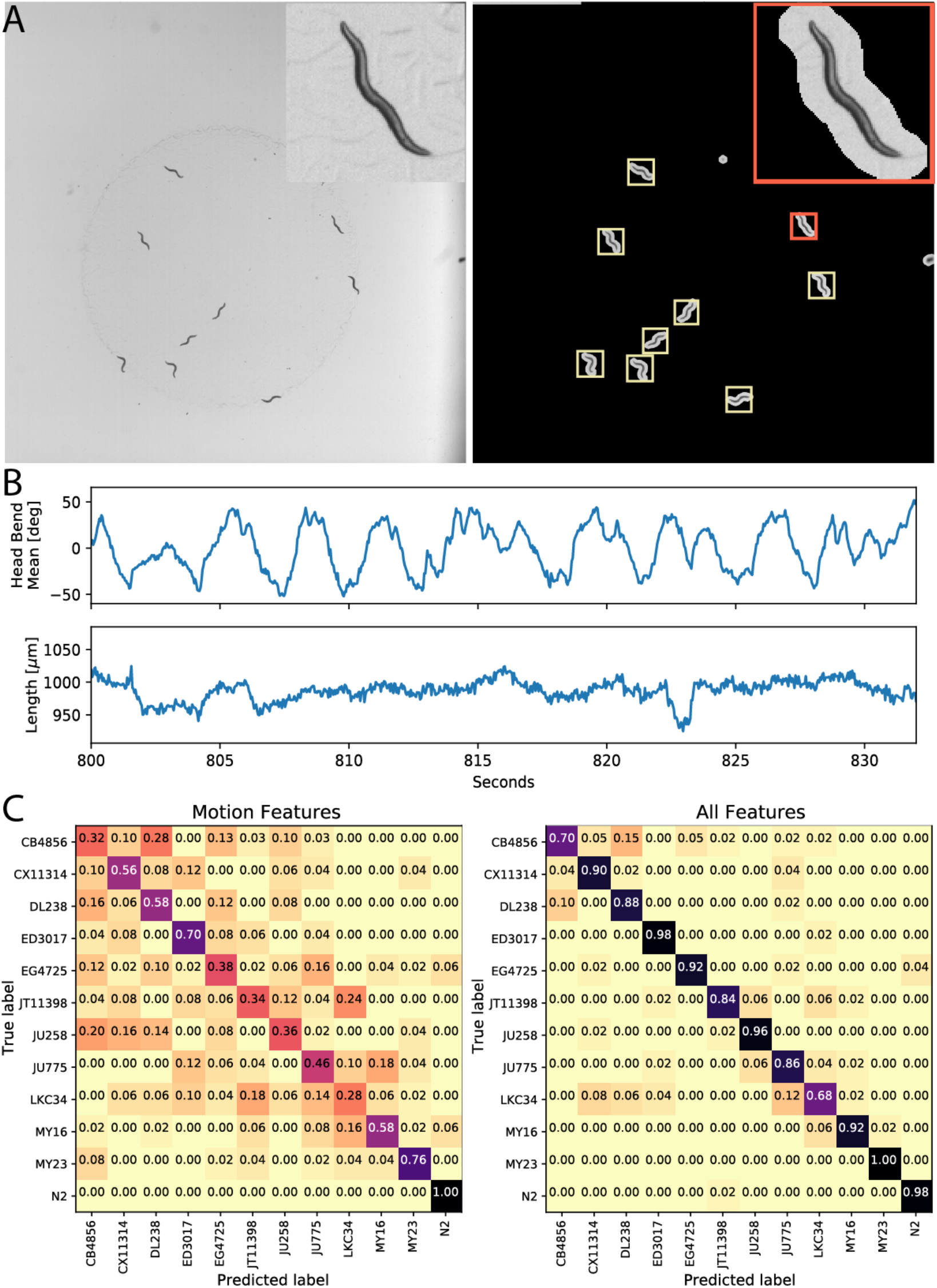
Classification of wild isolates. A. Left, original image. Right, results after the applying the compression mask. Each square corresponds to an identified worm. The worm in the red square is shown at a higher zoom in the inlets in each panel. B. Example of time series of two selected postural features. C. Confusion matrix of the classifier using 10-fold cross validation on features related with motion that could be extracted using low resolution multi-worm tracking (*left*) or using all the features we calculate using both motion and postural data (*right*).

**Fig. 6:**
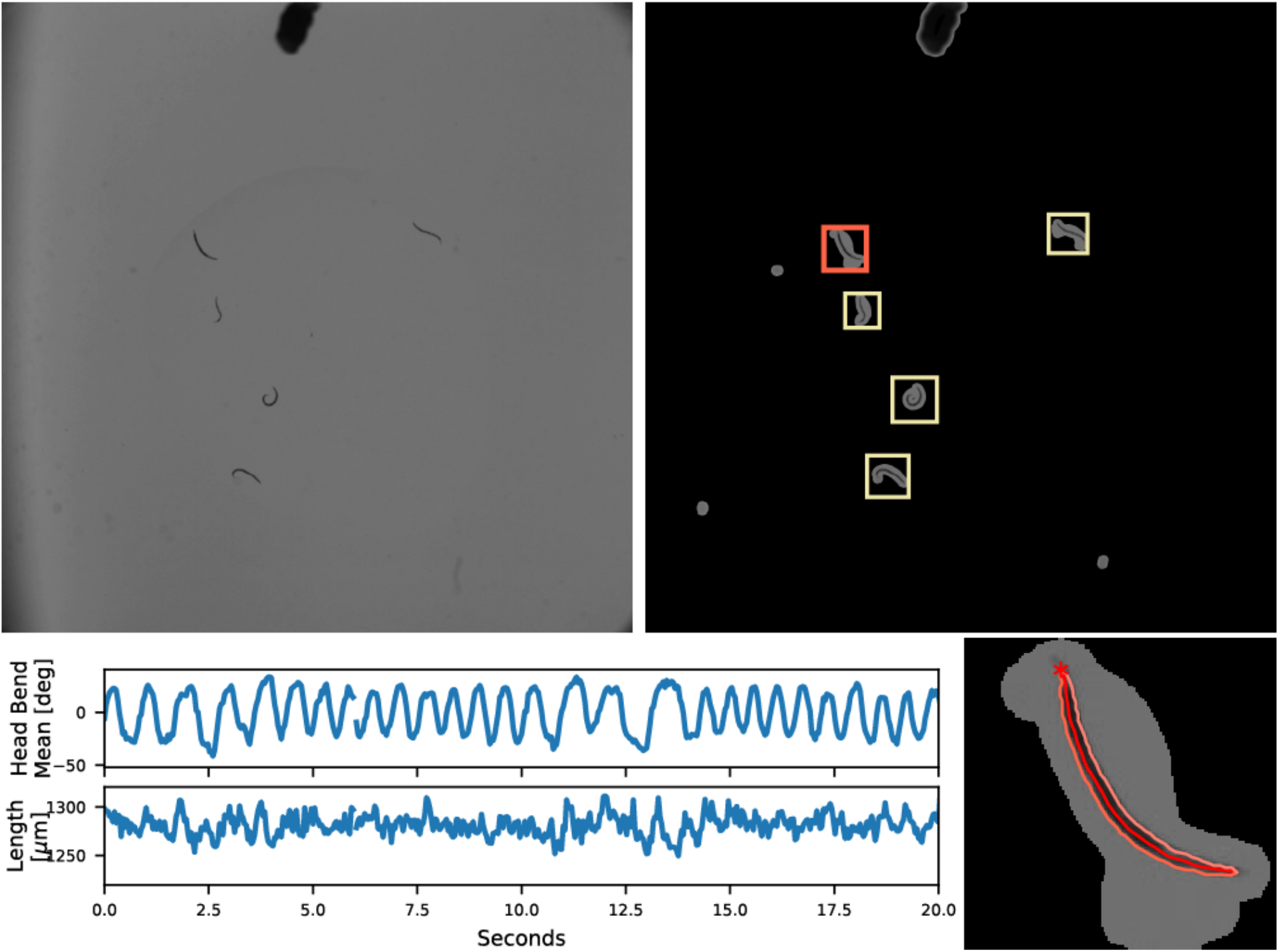
Swimming *C. elegans*. *Top left*, original image. *Top right*, results after applying the compression mask. Each square corresponds to an identified worm. *Bottom left*, example time series of two selected postural features. *Bottom right*, skeleton and contour of the worm inside the red square in the top right panel. The head is indicated with the red asterisk.

**Fig. 7:**
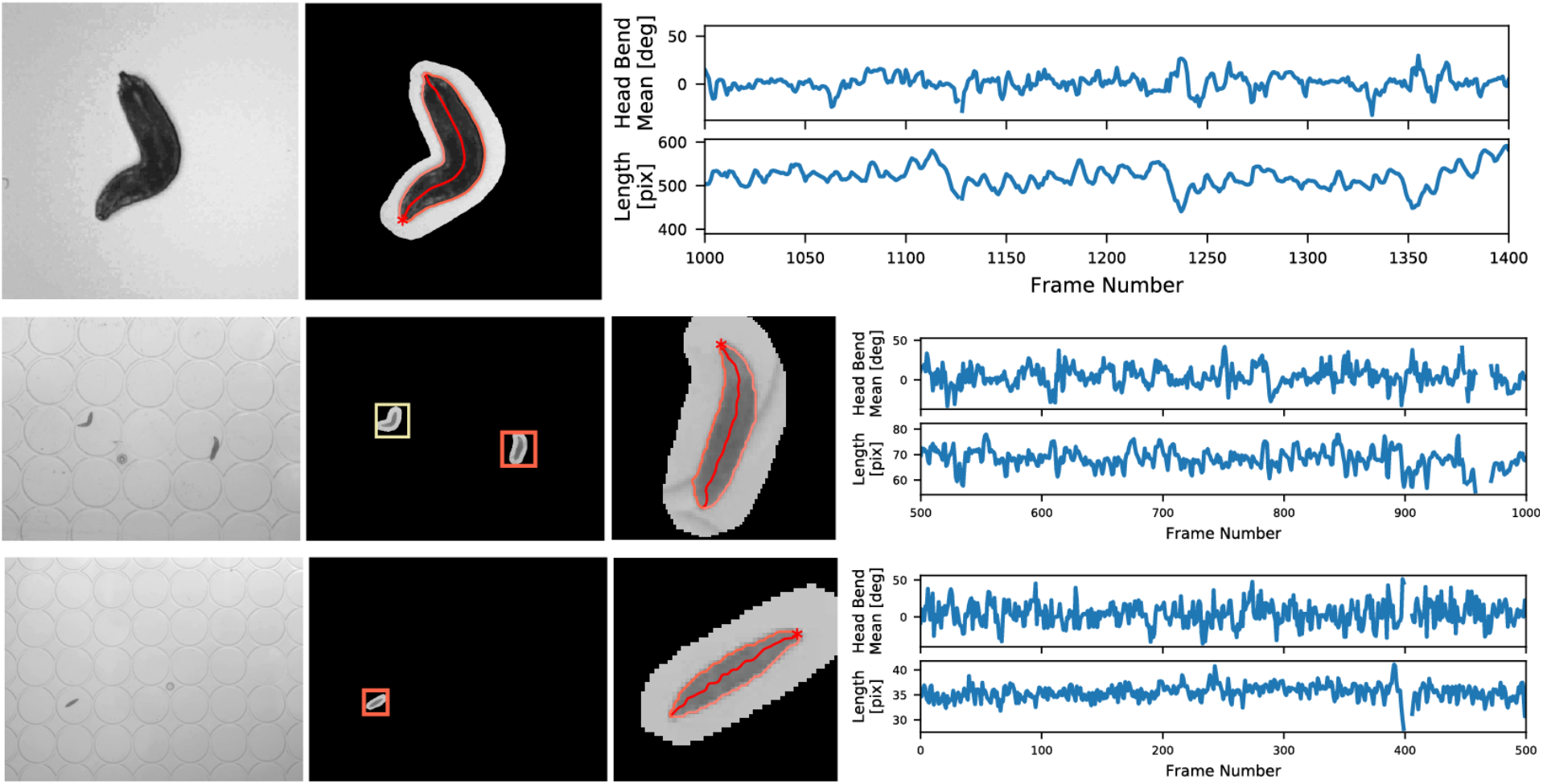
Videos of maggots at different resolutions. Data from ref^27^ (*top row*) and ref^28^ (*centre and bottom row*). The raw videos generously provided by Alex Gomez-Marin. Data from left to right: original image; results after applying the compression mask (each square corresponds to an identified maggot); skeleton and contour of a selected maggot; example time series of two selected postural features.

**Fig. 8:**
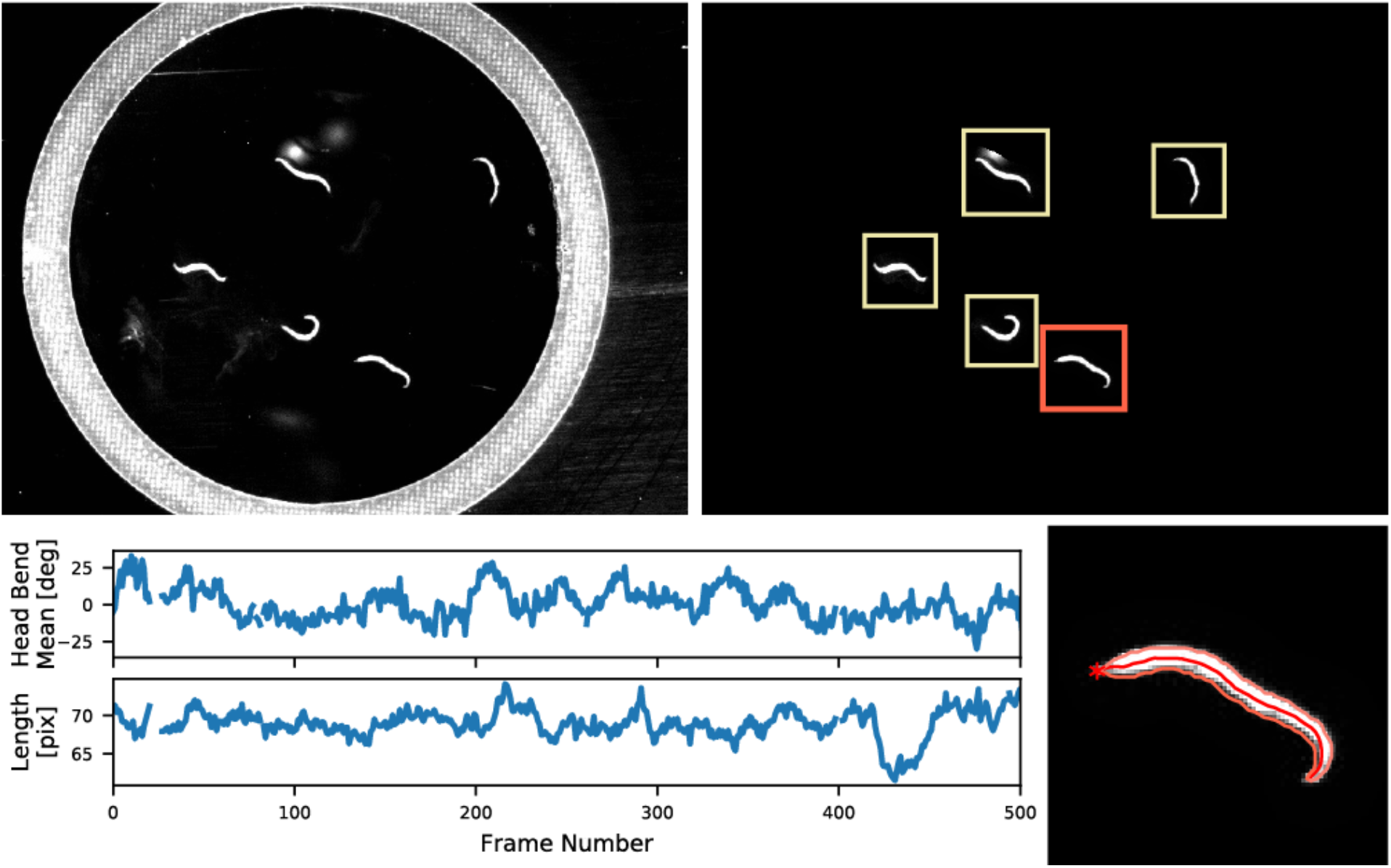
Dataset S1 in Restif *et al*.^6^. *Top left*, original image. *Top right*, results after the applying the compression mask. Each square corresponds to an identified worm. *Bottom left*, example time series of two selected postural features. *Bottom right*, skeleton and contour of the worm inside the red square in the top right panel. The head is indicated with the red asterisk.

**Fig. 9:**
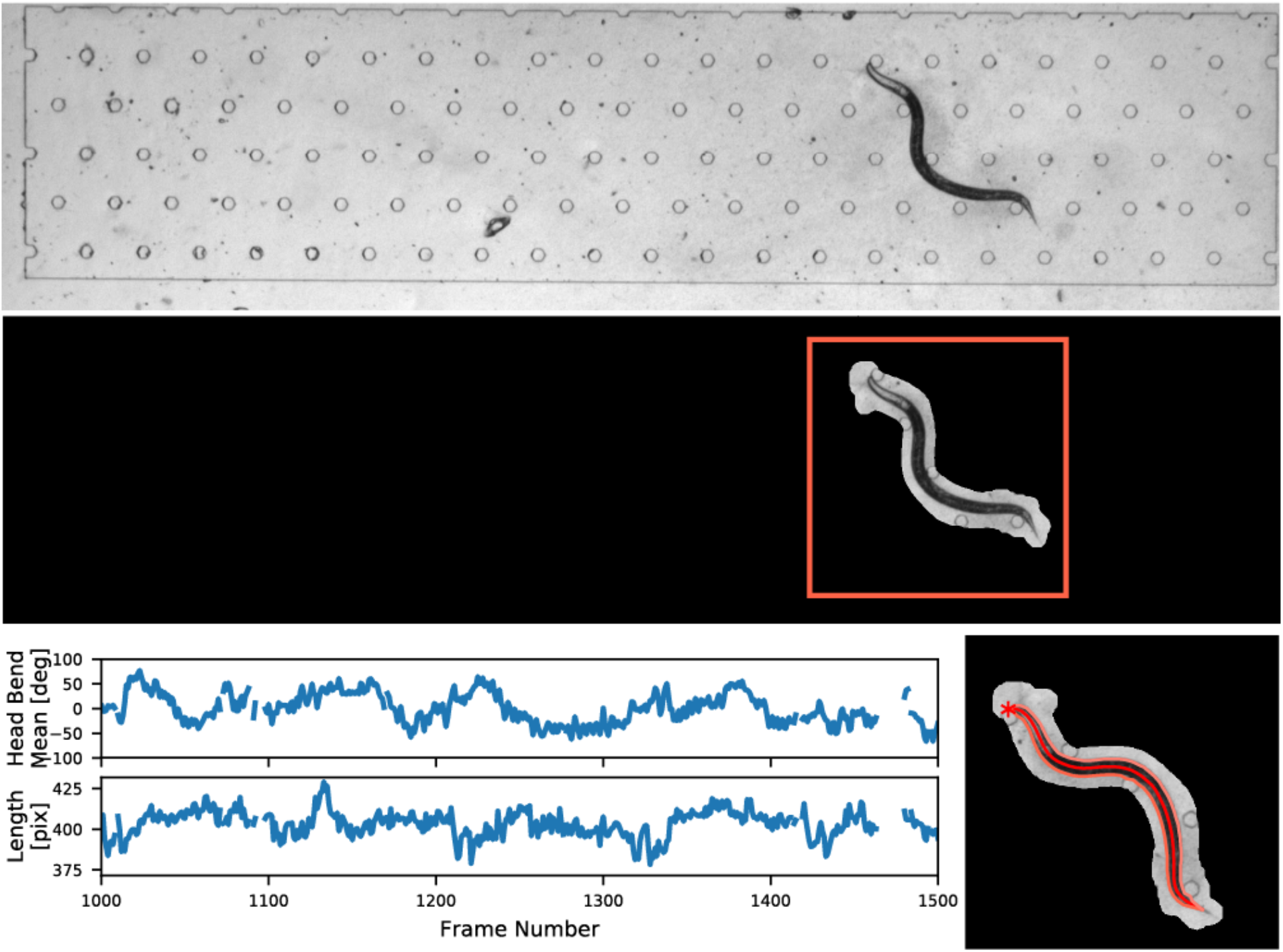
Example from pycelegans-1.0^7^. Data obtained from: *https://github.com/david-biron/pycelegans-1.0/tree/master/example/input*. *Top*, original image. *Middle*, results after the applying the compression mask. Each square corresponds to an identified worm. *Bottom left*, example time series of two selected postural features. *Bottom right*, skeleton and contour of the worm inside the red square in the middle panel. The head is indicated with the red asterisk.

**Fig. 10:**
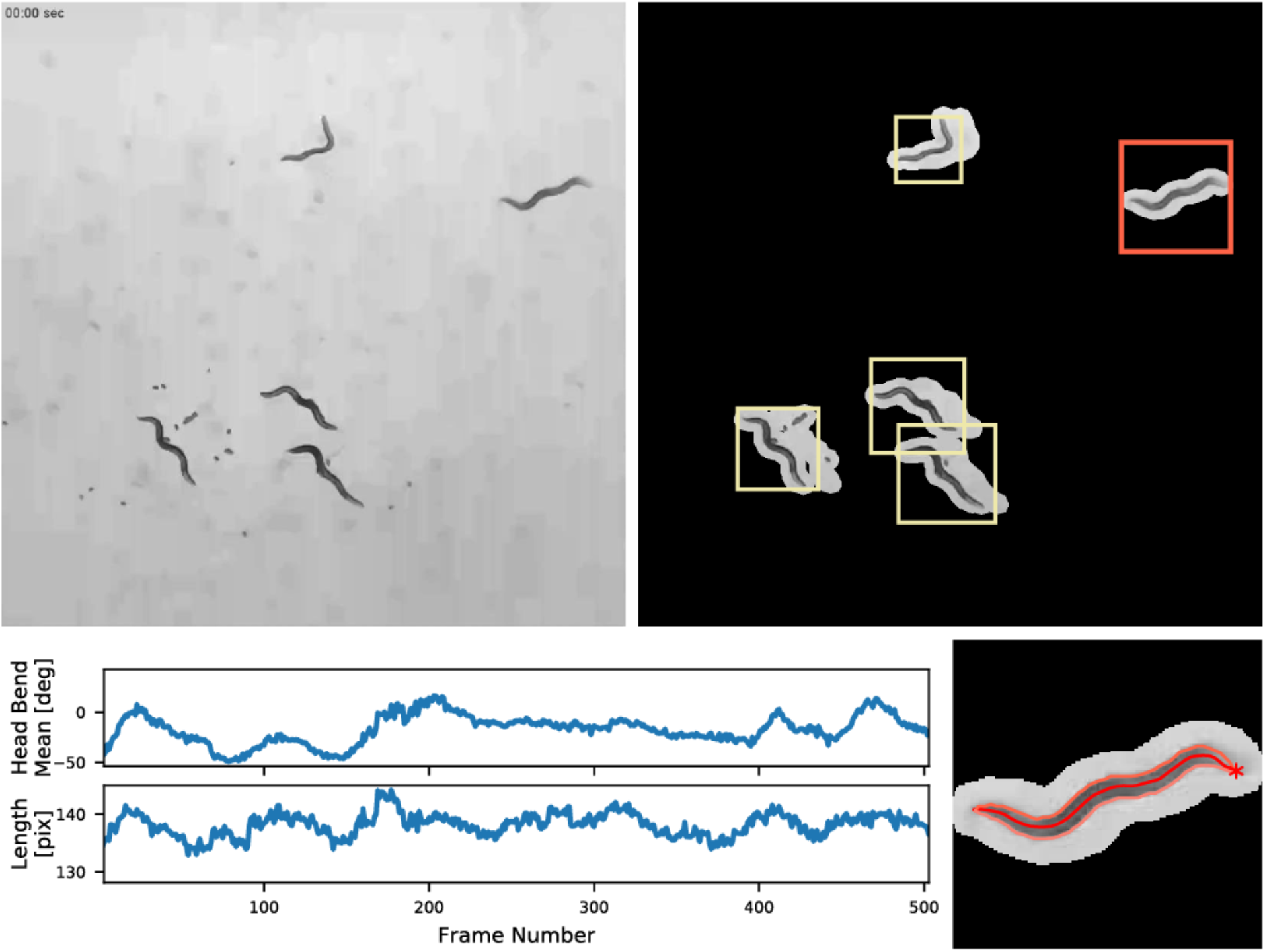
Video S6 in Chagas *et al*.^29^. *Top left*, original image. *Top right*, results after the applying the compression mask. Each square corresponds to an identified worm. *Bottom left*, example time series of two selected postural features. *Bottom right*, skeleton and contour of the worm inside the red square in the top right panel. The head is indicated with the red asterisk.

Although segmentation followed by lossless compression is our currently favoured approach, Tierpsy Tracker can use video codecs supported by the FFmpeg library (https://www.ffmpeg.org/) via OpenCV (https://opencv.org/), as well as setups where each frame is saved as an individual image. Tierpsy Tracker also makes it straightforward to save only worm contour, skeleton, and feature data because it creates a parallel directory structure for analysis results. If tracking results are acceptable, the original and/or compressed videos can simply be deleted to mimic the online behaviour of the Multi-Worm Tracker.

### Compression/object detection algorithm

The aim of the initial segmentation is to identify regions in the image that are likely to contain worms and to compress the data by only keeping the pixels around these candidate regions. The worms are assumed to be dark objects on a light background or light objects on a dark background. The detection algorithm uses an adaptive mean thresholding to account for local variations of the image intensity, filters the resulting blobs by their size, and uses morphological operations to clean the resulting mask and save the surrounding pixels. Each frame is then masked and saved into an HDF5 file using a compressed 3D array (frame * image width * image height). The HDF5 format is used due to its capacity to deal with arbitrarily large files, built-in compression filters, and its support of a large variety of software platforms.

Instead of calculating a new mask for each frame, we can save computation time by collecting a stack of frames and calculating the mask on the z-projection (minimum for a light background and maximum for a dark background). Using this approach together with the faster lz4 compression, a C++ implementation of the algorithm was implemented in our setup to run in real time (25 fps).

### Head/tail identification algorithm

The output of the skeletonization algorithm does not determine which end of the curve is the head and which one the tail. To orient the skeletons, we use the following algorithm which depends in part on having the high-resolution textural information in Tierpsy’s HDF5 file format:

1. During the skeletonization step we attempt to keep the same orientation between consecutive skeletons by choosing the orientation that minimizes the L2-norm of the current and previous skeletons. For this step, the orientation does not have to be correct (head and tail could be misplaced), we are only interested in keeping a spatial coherence between consecutive frames. This approach will maintain the same
2. orientation even if the skeletonization fails for few frames, as long as the time gap between consecutive skeletons is small relatively to the worm displacement.
3. We divided the movie into blocks of skeletons with the same orientation. A skeleton is assigned to a given block if there is not a gap larger than 0.5s with the previous skeleton in the block. For each block, we estimate the motility of each extreme in the skeleton as the rolling standard deviation (window 5s) of the angles between each end-point and a point at ~1/10 of the worm length from it. The end corresponding to the head is assumed to have a larger motility than the end corresponding to the tail. This step typically orients most of the skeletons correctly, but it will fail if the tail moves more in that particular block, which occurs most often in small blocks.
4. To correct for mistakes in the previous step we make use of the pixel intensities along the worms. We interpolate along the segments perpendicular to the worm skeletons to obtain the profile intensity along a straightened worm^25^ (Fig 4A). If the trajectories are large enough most of the skeletons in the previous step should have been oriented correctly, therefore if we get the median value of all the intensities we can determine a global intensity profile for each worm with the correct orientation (Fig 4B). Blocks of skeletons with incorrect orientation will show a switched intensity profile (Fig 4C). We detect these blocks by calculating the L1-norm of the difference between each frame profile and the global profile in both the correct orientation and with a switched orientation. If the L1-norm of the switched profile is smaller than that of the correct orientation, it is likely that the corresponding skeleton needs to be corrected.
5. The previous step will typically orient all the skeletons with the same orientation, but this orientation could be wrong in rare cases where most of the head/tail blocks were assigned incorrectly in step 2). As a final refinement step, we recalculate the motility of each extreme in the skeleton similar to 2) but using the whole video as a single block. In cases where the worm is mostly immobile, *i.e*. the total head displacement range is less than half the worm length, we increase the rolling window from 5s to 250s in order to increase the sensitivity of the analysis.

### Sample analysis

To demonstrate the usefulness of postural features in a multi-worm tracker, we used Tierpsy Tracker to extract features from videos of twelve wild isolated strains of *C. elegans* (the CeNDR divergent set^26^) and trained a classifier to distinguish them. We collected between 25 and 28 videos per strain. Each video contains 5-10 worms per plate. We pooled the time series and event data of all the worms in a given video and calculated the 726 morphological and behavioural features described in Yemini *et al*.^20^ We remove features where more than 5% of the videos have NaN values leaving 681 features. We imputed any remaining NaN values with the global average (i.e. the average of that feature across all strains) and z-normalised each feature by subtracting the mean and dividing by its standard deviation.

To classify the strains, we implemented a multinomial logistic regression as a Softmax layer using PyTorch. We trained for 500 epochs using a mini-batch size of 50 and stochastic gradient descent with learning rate 0.001 and momentum 0.9. Using a 6-fold stratified cross validation we obtained an accuracy 87.8 ± 4.0% (mean ± standard deviation) using all the 681 features selected as described above. To illustrate the importance of extracting high-resolution postural information, we trained a separate classifier on the same data using only features that could be extracted using a lower resolution multi-worm tracker without detailed postural data. These features are derivate from the midbody speed (linear and angular) and crawling (frequency and amplitude), motion events (*e.g*. turning rate, forward motion time, average displacement when moving backwards), and the path range and curvature. We selected in total 133 features and obtain a lower accuracy of 57.6 ± 4.5% (mean ± standard deviation).

Sample preparation: The protocol for tracking is similar to that described in Yemini *et al*. (2013). L4 larvae are picked onto a plate with OP50 and allowed to grow overnight to adulthood. Before imaging, 5 or 10 young adults are picked onto a 35 mm plate and allowed to habituate for 30 minutes before recording. Each recording lasts for 15 minutes. Imaging plates contain nematode growth medium (NGM) with low peptone that have been seeded the day before 75μL of OP50.

### Examples of Tierpsy Tracker analysis using different experimental setups

A useful characteristic of Tierpsy Tracker is that it can deal with data from a large variety of experimental setups. We provide the following examples using previously published data:

- Fig. 6 Swimming *C. elegans*.
- Fig. 7 *Drosophila* larvae^27,28^.
- Fig. 8 Dataset S1 in Restif *et al*.^6^
- Fig. 9 Example from pycelegans-1.0^7^.
- Fig. 10 Video S6 in Chagas *et al*.^29^

